# Shell shape as an indicator of phenotypic stocks of Tehuelche scallop (*Aequipecten tehuelchus*) in Northern Patagonia, Argentina

**DOI:** 10.1101/2021.10.14.464278

**Authors:** Leandro Nicolás Getino Mamet, Gaspar Soria, Laura Schejter, Federico Márquez

## Abstract

Tehuelche scallop, *Aequipecten tehuelchus*, is a commercially exploited species in Northern Patagonia, Argentina. Without genetic differentiation at the species level, *A. tehuelchus* presents three morphotypes: *tehuelchus*, *madrynensis*, and a non-common variant *felipponei*. The main goal of this study was to analyze the shell shape variation of Tehuelche scallop to differentiate and identify the phenotypic stocks. The shape differences between and within the two main morphotypes (*tehuelchus* and *madrynensis*) were assessed using geometric morphometrics in nine localities. The shell shape presented variability at geographic scale, with the morphologic traits that maximized the differentiation among localities between the *tehuelchus* and *madrynensis* morphotypes. Scallops from *madrynensis* morphotype presented higher and circular shell discs with smaller auricles than those from *tehuelchus* morphotype. Morphometric differentiation was also detected among localities of each morphotype, wherein most of the variability was related to the disc circularity and the symmetry of the auricles. The presence of morphologic variation in San Matías and San José gulfs, wherein a single genetic pool is shared, evidenced the plastic nature of the species. Given the distribution of this resource in distinct provincial jurisdictions, the differentiation of phenotypic stocks has relevance in the context of fishery management, especially if zoning and rotational strategies are implemented.

## 1. Introduction

Phenotypic plasticity refers to the alternative phenotypes (morphology, physiology, behavior, or other environmental sensitive traits) that a given genotype responds to different ecological conditions (DeWitt and Scheiner, 2004; Nicolaus and Edelaar, 2018; Schlichting and Pigliucci, 1998; Sommer, 2020). This flexible response could be relevant to cope with the environmental heterogeneity, being especially relevant in sessile and sedentary organisms that cannot escape or avoid changes in environmental conditions (Nicolaus and Edelaar, 2018). Particularly, the shell morphology of marine mollusk species is recognized for being susceptible to environmental conditions (Márquez et al., 2017a; Melatunan et al., 2013; Peyer et al., 2010; Scherer et al., 2016; Urdy et al., 2010).

In marine shell-bearing invertebrates, shell shape variation is relevant to limit phenotypic stocks (*sensu* Booke, 1981), as were studied for distinct molluscan species with commercial interest (i.e., Márquez et al., 2017b, 2010b; Palmer et al., 2004; Rufino et al., 2013; Trivellini et al., 2021). The use of geometric morphometric techniques has been recognized to help identify phenotypic stocks of marine mollusks (Cadrin, 2020; Márquez et al., 2017b, 2010b; Palmer et al., 2004; Rufino et al., 2013). The identification of stocks is especially relevant for spatially structured populations to define spatial management tools (e.g., no-take zones, zoning, and rotation schemes) or to identify the origin of the landings (Cadrin, 2020; Ibáñez, 2015).

Tehuelche scallop, *Aequipecten tehuelchus* (d’Orbigny, 1846) is one of the main coastal shellfish resources that sustain a small-scale fishery in Northern Patagonia, Argentina (Elías et al., 2009; Getino Mamet et al., 2021). The species is found in shallow waters of the Southwestern Atlantic Ocean, between 23° to 45° latitude S, along the Argentine and Magellanic biogeographic provinces (Fig. 1). At the southern end of its distribution range, the presence of oceanographic fronts divides water masses in distinct regions (Acha et al., 2004). Such fronts include El Rincón haline front between 39°S–41°S (Acha et al., 2004), the San Matías Gulf thermal front (Tonini et al., 2013), the Peninsula Valdes tidal front (Pisoni et al., 2015), and the San José Gulf tidal front (Amoroso and Gagliardini, 2010) (Fig. 1). In particular, the San José Gulf tidal front divides the homonym gulf in two hydrographic domains (east and west) that define the local demography of Tehuelche scallop (Amoroso et al., 2011; Amoroso and Gagliardini, 2010).

**Figure 1.**
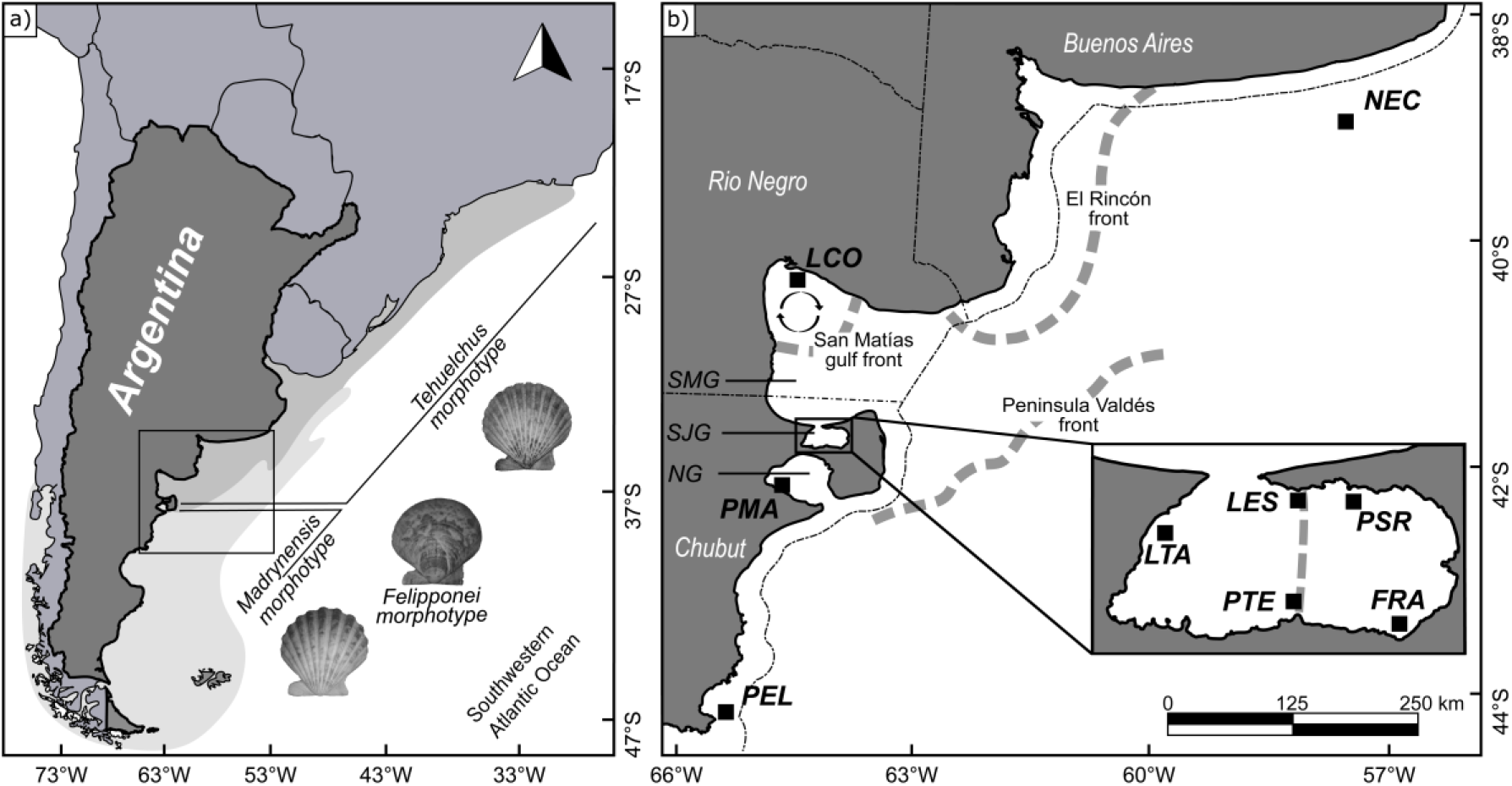
Study area. a) Distribution of *Aequipecten tehuelchus* depicting the *tehuelchus*, *madrynensis*, and *felipponei* morphotypes. The shaded region indicates the location of the Argentine (northern) and Magellanic (southern) biogeographic provinces. b) Sampling locations: Necochea (NEC), Las Conchillas (LCO), La Esfinge (LES), Punta San Román (PSR), Fracasso (FRA), Punta Tehuelches (PTE), La Tapera (LTA), Puerto Madryn (PMA), and Playa Elola (PEL). The Northern Patagonian gulfs which encompass San Matías, San José, and Nuevo gulfs are indicated (SMG, SJG, NG respectively). Gray dashed lines indicate the location of El Rincón front, San Matías Gulf front and associated cyclonic gyre, San José Gulf tidal front, and Peninsula Valdes front. The dot-dashed lines mark political borders between Buenos Aires, Río Negro, and Chubut provinces, and, to the east, the limit between provincial and federal jurisdictions over the Argentine Sea.

The species present three morphotypes: *tehuelchus, madrynensis*, and a non-common variant *felipponei*. The *tehuelchus* and *madrynensis* morphotypes are geographically separated in a distribution that matches the Argentine and the Magellanic Biogeographic provinces, respectively (Castellanos, 1971; Soria et al., 2016) (Fig. 1a); meanwhile, *felipponei* is scarcely found in both biogeographic provinces (Schejter and Bremec, 2007; Trovant et al., 2019). Those have been meristically differentiated by the number of ribs as follows: *tehuelchus* has 14–19 ribs, *madrynensis* has fewer ribs (11–14) but more pronounced, and *felipponei* has no ribs (Castellanos, 1971). Genetic studies using allozymes (Real et al., 2004) and mitochondrial markers (Trovant et al., 2019) indicated an absence of differentiation among morphotypes at the species level. On the other hand, employing more variable microsatellites markers, genetic differentiation at the population level was detected between *tehuelchus* and *madrynensis* morphotypes following the biogeographical distribution of the morphotypes (Getino Mamet et al., 2021). Within the *tehuelchus* morphotype, scallops also present allometric variations (Ciocco, 1992; Márquez et al., 2010a). Juvenile scallops present circular shells with symmetrical auricles, while adults present elliptical shells and smaller auricles (Márquez et al., 2010a).

The Tehuelche scallop fishery of San Matías and San José gulfs take place, respectively, in Río Negro and Chubut provincial jurisdictions that are managed independently. The fishery management in each provincial jurisdiction differs in the fishing method, the access regime, stock assessment, and quota allocation (Orensanz et al., 2007; Orensanz and Seijo, 2013, Getino et al., 2021). However, these scallop demes, belong to a single genetic stock being part of the *tehuelchus* morphotype (Getino Mamet et al., 2021). Combining captures from San José and San Matías gulfs, reported landings in 2020 were less than a thousand tons (Getino Mamet et al., 2021). On the other hand, northerly and outside of this regular fishing region, a Tehuelche scallop bed was incidentally detected and exploited in a single fishing pulse near Buenos Aires Province in 2002 by the industrial Patagonian scallop fleet (Lasta and Campodónico, 2003; Soria et al., 2016) without fishing records afterward. The absence of scallop beds in Nuevo Gulf and further south precluded the development of any kind of fishing activity at any scale.

In this context, the main goal of this study was to describe the shell shape differences of Tehuelche scallop among nine locations to differentiate and identify the phenotypic stocks throughout the distributional range of the species within the Argentine Sea with particular interest where the fishery takes place. To accomplish this, the shape differences between and within *tehuelchus* and *madrynensis* morphotypes were assessed using geometric morphometric techniques (Adams et al., 2004).

## 2. Materials and methods

A total of 307 Tehuelche scallop shells were collected in nine localities: Necochea (NEC), Las Conchillas (LCO), La Esfinge (LES), Punta San Román (PSR), Fracasso (FRA), Punta Tehuelches (PTE), La Tapera (LTA), Puerto Madryn (PMA), and Playa Elola (PEL), (Fig. 1b, Table 1). The outline of the interior size of the right valve of each scallop was digitized using a scanner EPSON Perfection V370 Photo at 600 PPI.

**Table 1.** Sample locations. Sampling locations indicating the code; morphotype; status of the fishery (fishing gear between brackets as following: BT, bottom trawls; AD, artisanal dredge; and HO, hookah-diving); latitude; longitude; sample size (N); and year of collection.

| Location | Code | Morphotype | Status of fishery | Latitude | Longitude | N | Year |
| --- | --- | --- | --- | --- | --- | --- | --- |
| Necochea | NEC | *tehuelchus* | Single pulse (BT) | 39°30'S | 59°30'W | 54 | 2002 |
| Las Conchillas | LCO | *tehuelchus* | Resumed activity* (AD/HO) | 40°53'S | 64°45'W | 51 | 2017 |
| La Esfinge | LES | *tehuelchus* | Regular activity (HO) | 42°15'S | 64°17'W | 30 | 2012 |
| Punta San Román | PSR | *tehuelchus* | Regular activity (HO) | 42°15'S | 64°13'W | 33 | 2012 |
| Fracasso | FRA | *tehuelchus* | Regular activity (HO) | 42°24'S | 64°08'W | 24 | 2012 |
| Punta Tehuelche | PTE | *tehuelchus* | Regular activity (HO) | 42°24'S | 64°19'W | 24 | 2012 |
| La Tapera | LTA | *tehuelchus* | Regular activity (HO) | 42°18'S | 64°32'W | 26 | 2012 |
| Puerto Madryn | PMA | *madrynensis* | Fishing is absent | 42°46'S | 65°00'W | 37 | 2017 |
| Playa Elola | PEL | *madrynensis* | Fishing is absent | 44°53'S | 65°38'W | 28 | 2016 |

\*After fifteen years of absence of the fishery, a new recruitment pulse was recorded in 2019-2020 in the NE sector of the San Matías Gulf and a dredge-based fishery is currently active under the Rio Negro Provincial Jurisdiction.

The analysis of the shell shape was performed using landmark-based (2D) geometric morphometric techniques, which allow preserving the shape information during the multivariate analyses so results can be graphically displayed (Adams et al., 2004). The shell outline was captured using a configuration with six landmarks (LM) and 23 semi-landmarks (S-LM) (Fig. 2). All specimens were digitized by the same observer using tpsDig2 v.2.17 (TPS series, available at https://life.bio.sunysb.edu/morph/index.html). The semi-landmarks were used to capture contours between LM, which were homologated mathematically in an iterative process (sliding) using TpsRelw v.1.53. In this method, the S-LM coordinates are slide along the outline profile of the scallop to minimize the bending energy of the landmark configuration (Gunz et al., 2005). Subsequently, a generalized Procrustes analysis was applied in which landmark configurations were rotated, translated to a common origin, and scaled to a unitary centroid size to obtain the Procrustes aligned coordinates, used as shape data (Bookstein, 1996).

**Figure 2.**
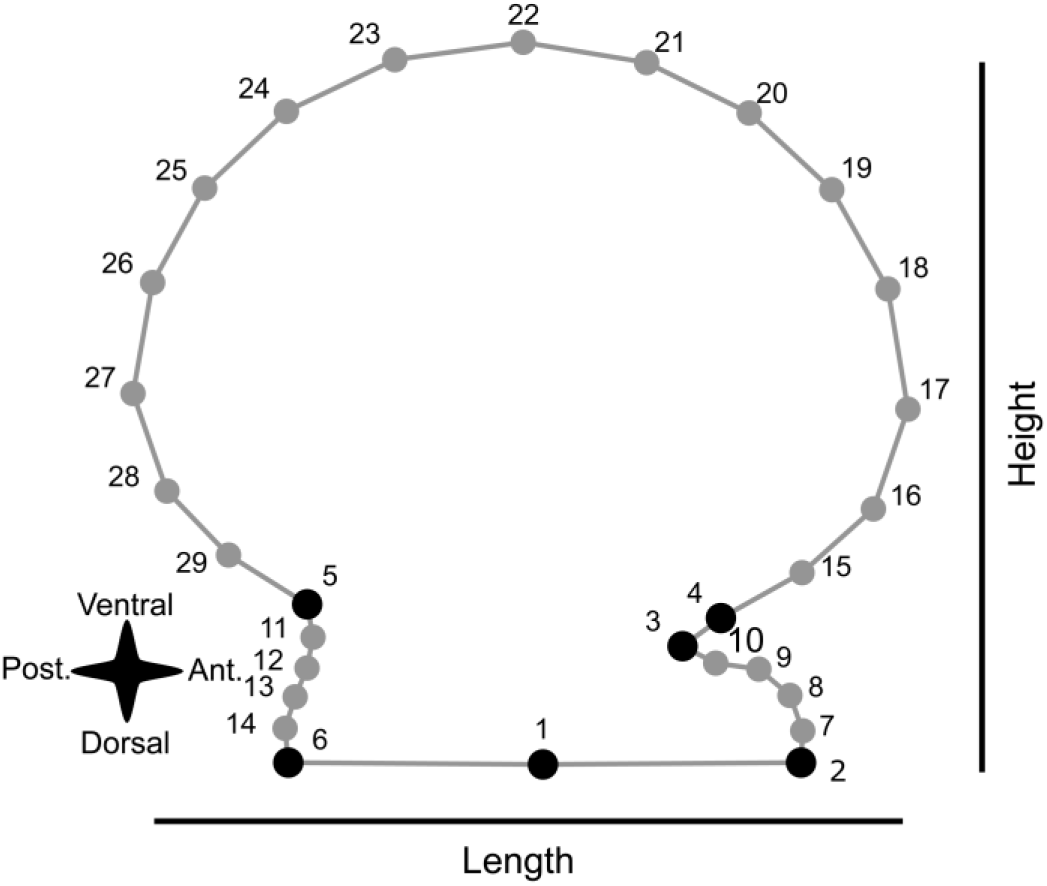
Shell shape analysis. Position of six landmarks (black dots) and 23 semi-landmarks (grey dots) used in the geometric morphometric analysis. Numbers refer to each landmark definition: LM 1, the base of hinge; LM 2, end of the anterior auricle; LM 3, posterior end of the byssal notch; LM 4, anterior end of the byssal notch; LM 5, inflection point between the shell disc and posterior auricle; LM 6 end of the posterior auricle; S-LM 7-10, equidistantly between LM 2 and LM 3; S-LM 11-14, equidistantly between LM 5 and LM 6; and S-LM 15-29, equidistantly between LM 4 and LM 5 on the shell disc. Shell orientation (anterior, Ant.; and posterior, Post.) and basic dimension are indicated.

A multivariate regression was performed in MorphoJ v1.06b (Klingenberg, 2011) in order to account for the shape changes related to the individuals’ size and discard this source of variation from the spatial analyses. Therefore, further analyses were made using the residual variation of the multivariate regression as free-size shape variables (Zelditch et al., 2004). The use of such variables can prevent the misinterpretation of among-localities differences if they are correlated to size (Outomuro and Johansson, 2017). The multivariate regression (pooled within localities) was made using the centroid size (CS), defined as the square root of the sum of the squared distances from the LMs to the centroid which they define, as shell size (Zelditch et al., 2004). Procrustes coordinates were used as shape variables. The significance was tested with a permutation test (100.000 rounds).

To elucidate the shape variation that maximizes the discrimination among localities, a canonical variate analysis (CVA) was performed in MorphoJ. An assignation table for each locality was constructed from the CVA using cross-validation. In the cross-validation, each individual is left out from the analysis and then assigned to one group (a locality in this case) using a function calculated from all other specimens (Kovarovic et al., 2011). This way avoids the biases caused by classifying one specimen using functions that were calculated on samples that included that same specimen (Viscosi and Cardini, 2011).

To evaluate the morphologic differentiation between *tehuelchus* and *madrynensis* morphotypes, a discriminant function analysis (DFA) was performed in MorphoJ and the shape differences were tested by the T-square Hotelling test with a permutation test (100.000 rounds). Cross-validation was also performed to check the accuracy of the discriminant function in assigning individuals to each group (morphotypes in this case).

Additionally, to summarize shape differences among localities, a hierarchical cluster analysis using an unweighted pair-group method with arithmetic mean (UPGMA) using Mahalanobis distances was performed in InfoStat. To identify the number of groups statistically significant, the UPGMA was coupled to an extension of a multiple comparison method based on cluster analysis (UPGMA-MDGC) (Valdano and Di Rienzo, 2007).

## 3. Results

The shell shape and CS were allometrically related (multivariate regression analysis, *P*<0.0001) and accounted for 11.1% of the total shape variation. The allometric component in shape was mainly explained by the shell circularity and the auricles symmetry. Larger individuals were represented by an elliptic shell disc, elongated in relation to the height, and shorter auricles. On the opposite, smaller individuals were represented by a more circular shell disc, and higher auricles (Fig. 3).

**Figure 3.**
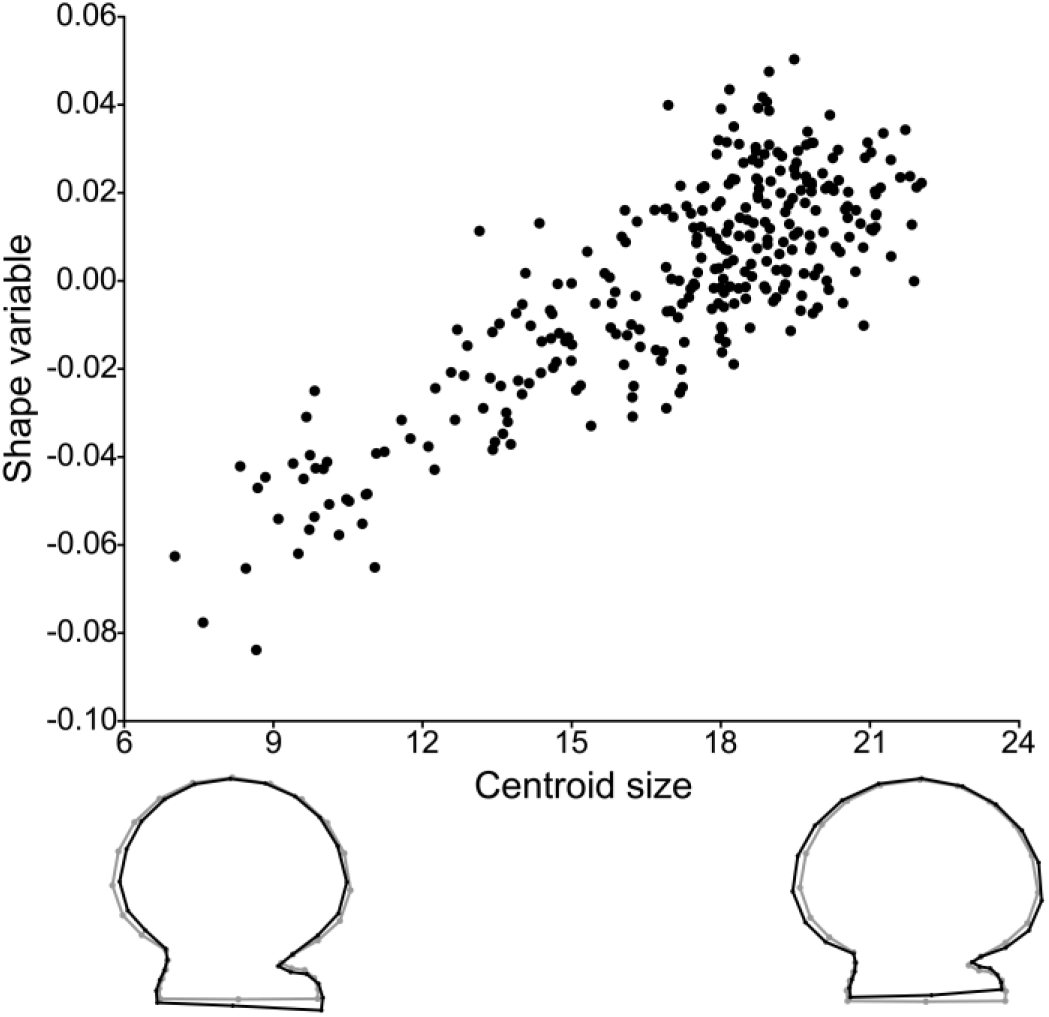
Allometric relationship between shape and size. The wireframe diagrams below the scatterplot, indicate the shape change predicted from consensus shape (grey wireframe) to an increment of ±10 scale factor of centroid size (black wireframe).

The shape variation that maximizes separation among localities is summarized by the CVA, in which the canonical variables 1 and 2 (CV1 and CV2, respectively) accumulated about 51.7% of the variation (Fig. 4). The shape components that differentiate the positive extreme of the CV1 were a higher and more circular shell disc and reduced auricles (Fig. 4). Tehuelche scallops from PEL were the most differentiated individuals, represented by positive values of CV1. Tehuelche scallops from both PMA and LES were also over the positive values of the CV1, although differenced over the CV2. On the other hand, the shell shape associated with the positive values of CV2 has a shorter height in relation to the length that results in an elliptically elongated shell-disc (in the anteroposterior sense) (Fig. 2), and symmetrically expanded auricles (Fig. 4). On the opposite extreme, the negative values of CV2 presented a shape characterized by a more circular and higher shell than the consensus and reduced and asymmetrical auricles. Scallops from LCO and PMA presented positive values of the CV2 wile PTE and LES were over the negative extreme of the axis (Fig. 4). Other locations, as PSR, FRA, or LTA, presented near-zero scores in both canonicals variables, while NEC was slightly displaced to the negative values of the CV2. The percentages of correct assignation were high in all locations, ranging between 82.3% (LCO) and 94.6% (PMA) (Table 2).

**Figure 4.**
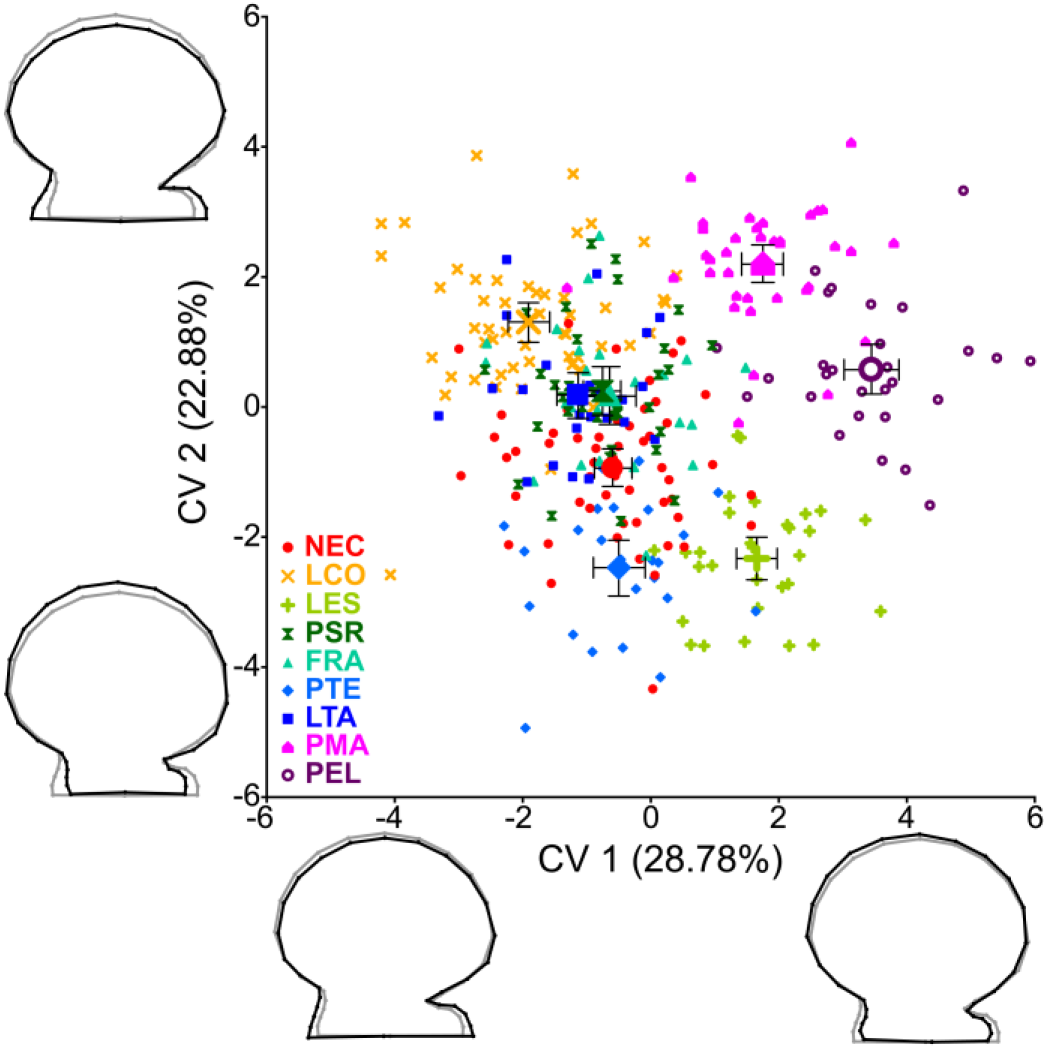
Analysis of the maximum variation in shell shape along the first two canonical roots. The expanded symbols indicate each location mean and whiskers are the 95% confidence interval for each mean. The wireframes diagrams show the deformation explained by each canonical variable axis with a scale factor of 10 to improve visualization. Grey color indicates the consensus shape, while black indicates the deformation over each canonical variable. Sampling locations: Necochea (NEC), Las Conchillas (LCO), La Esfinge (LES), Punta San Román (PSR), Fracasso (FRA), Punta Tehuelches (PTE), La Tapera (LTA), Puerto Madryn (PMA), and Playa Elola (PEL).

**Table 2.**
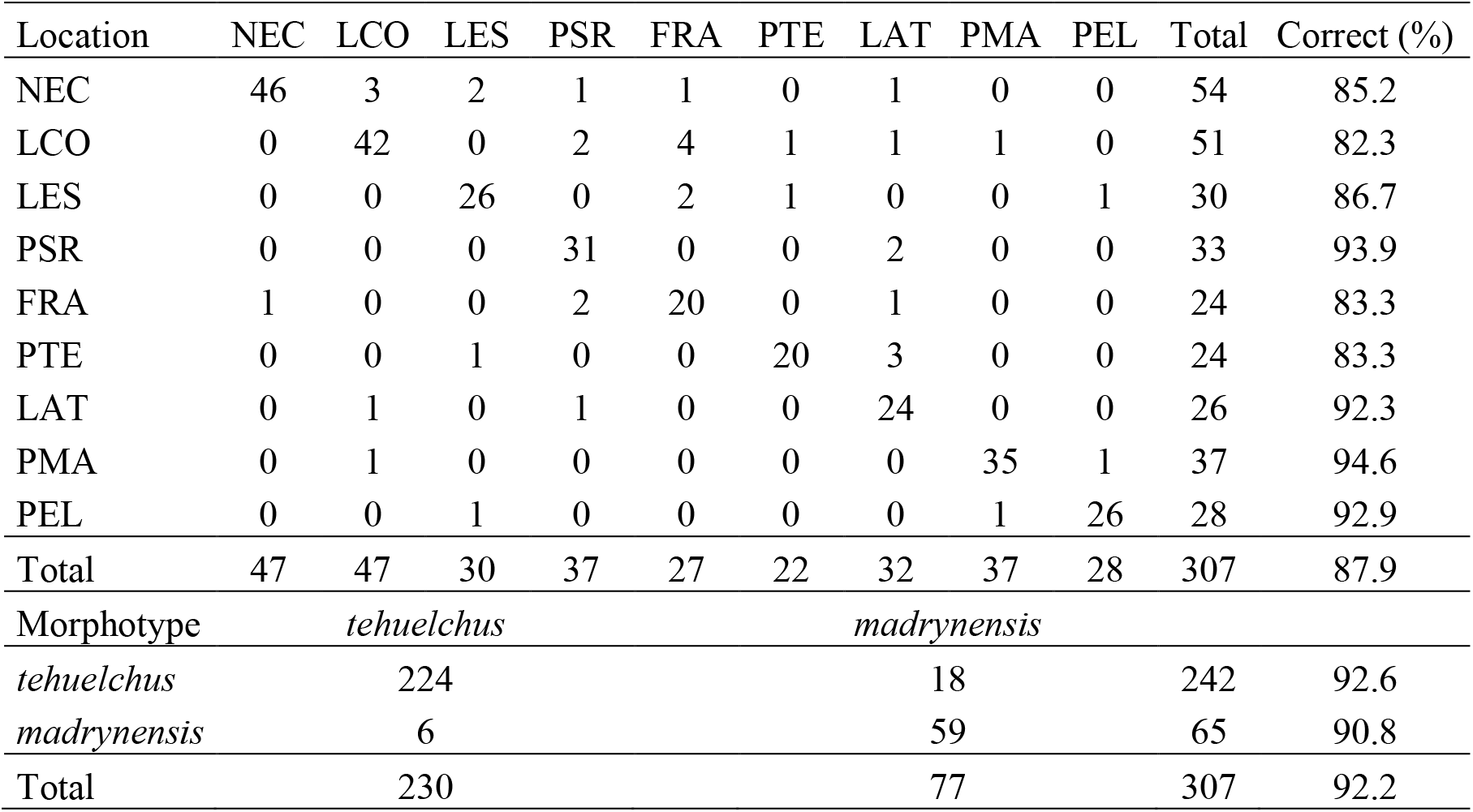
Assignation table for localities and morphotypes resulting from the cross-validation. Rows indicate the true group of origin (locality or morphotype), and columns indicate the groups in which individuals were allocated.

Shell shape between *tehuelchus* and *madrynensis* mophotypes differed significantly (DFA; *P* <0.0001). The maximum shell shape deformation that differentiates both morphotypes (Fig. 5) was similar to the explained by the first canonical variable of the CVA. The *madrynensis* morphotype presented a more circular shell with symmetric and reduced auricles; oppositely, the *tehuelchus* morphotype was associated with asymmetric auricles and an elongated shell outline (Fig. 5). The cross-validation of the discriminant function analysis assigned correctly 92.6% and 90.8% of *tehuelchus* and *madrynensis* samples, respectively (Table 2).

**Figure 5.**
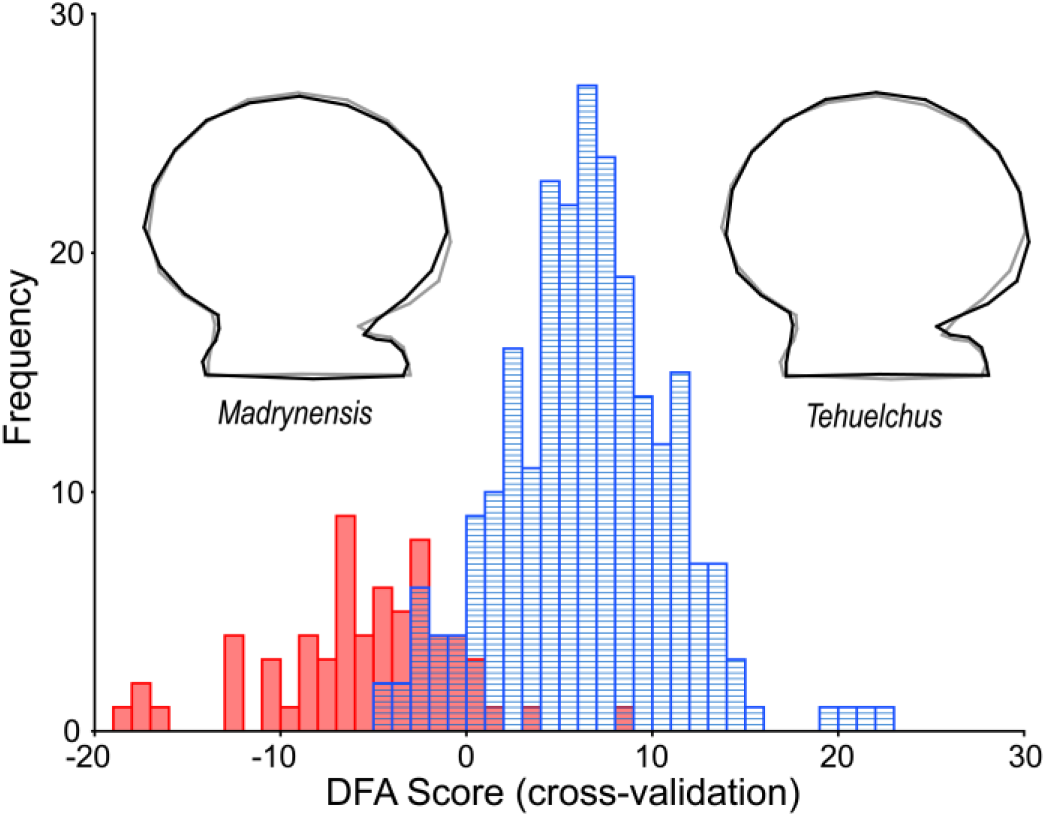
Discriminate function analysis (DFA) indicating the assignation frequency of individuals from each morphotype. The wireframes diagrams show the deformation associated with each morphotype, with a scale factor of 5 to improve visualization. Grey color indicates the consensus shape, while black indicates the deformation of each morphotype.

The pattern of location similarities (Mahalanobis distances), summarized by the UPGMA-MDGC dendrogram shows the main patterns of differentiation between morphotypes and clarified the similarities within *tehuelchus* locations. The most divergent group was compounded by PEL and PMA localities (*madrynensis* morphotype) (Fig. 6). Within the *tehuelchus* group, the most divergent shell shape was at PTE and LES. The remaining location formed an inner group wherein NEC was differenced from LCO, PSR, LTA, and FRA localities. Except for LTA and PSR which lack differentiation, the remaining locations present statistically differenced shapes.

**Figure 6.**
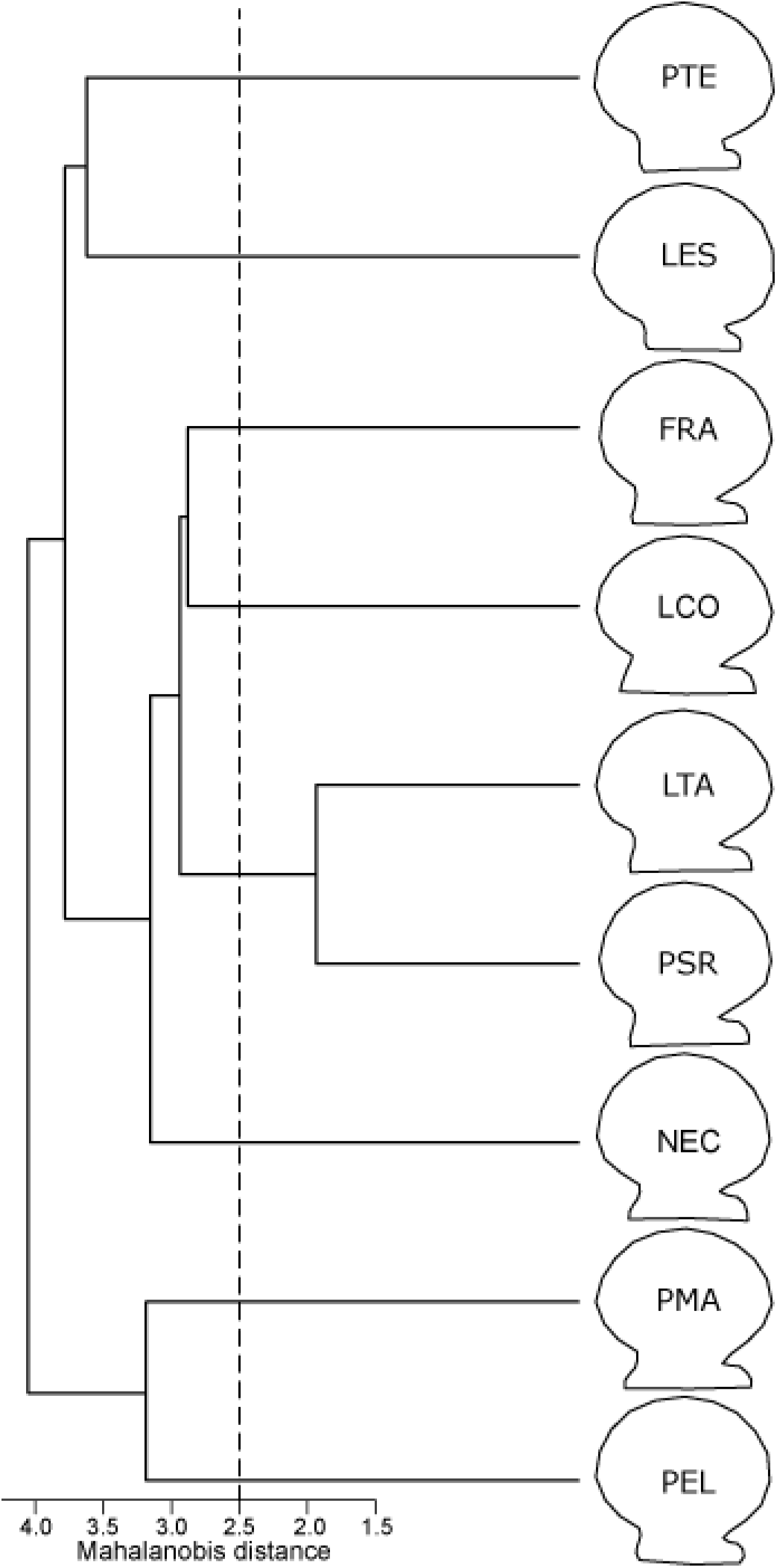
Dendrogram of the unweighted pair-group method with arithmetic mean (UPGMA) based on Mahalanobis distances. Branches indicate the patterns of shape similitude among locations. The vertical dashed line indicates the cut-off criterion for statistically distinct groups identified by the MDGC test (p=0.05). Shape diagrams represent the average shape for each location; deformation was exaggerated three times to improve the visualization. Sampling locations: Necochea (NEC), Las Conchillas (LCO), La Esfinge (LES), Punta San Román (PSR), Fracasso (FRA), Punta Tehuelches (PTE), La Tapera (LTA), Puerto Madryn (PMA), and Playa Elola (PEL).

## 4. Discussion

One of the most remarkable features of scallops is the swimming capability; such characteristic defines the main patterns of shell shape variation of these species (Stanley, 1988; Tremblay and Guderley, 2016). Additionally, the local environmental conditions prevailing where scallops live might be among the forces that drive the shell outline. In this context, Tehuelche scallop from northern Patagonia, Argentina, presents polyphenism wherein the shell shape variability of adults has been driven by oceanographic heterogeneity.

The principal variability of the scallops shell shape was detected between the *tehuelchus* and *madrynensis* morphotypes. The *madrynensis* morphotype presents higher and circular shell discs with reduced auricles than those from the *tehuelchus* morphotype. Also, significant differences were detected at PMA, with an elliptically elongated shell-disc (in the anteroposterior sense) and symmetrically expanded auricles. These findings complement the morphotype characterization defined solely based on the number of ribs (Castellanos, 1971; Real et al., 2004). The geographic limit between both morphotypes matches the boundary between the Argentine and the Magellanic Biogeographic provinces (Balech and Ehrlich, 2008; Cousseau et al., 2020). While the *tehuelchus* morphotype is found in the warm-temperate waters from the Argentine Biogeographic province, the *madrynensis* morphotype inhabits the cold-temperate Magellanic province (Real et al., 2004; Trovant et al., 2019). The presence of the Peninsula Valdés tidal front in this region (Pisoni et al., 2015) seems to act as a physical barrier that partially constrains larval dispersal (Alfaya et al., 2020; Getino Mamet et al., 2021). The isolation resulting from such barrier might be strong enough to produce both the morphologic and genetic differentiation between morphotypes (Getino Mamet et al., 2021). The findings of both genetic and morphologic differentiation within the species according to the oceanographic differentiation between biogeographic provinces could be indicative of the presence of two ecotypes.

On the other hand, high shell variability was found among locations of the northern *tehuelchus* morphotype. Most of the variability is related to the symmetry of the auricles and the shell disc circularity, the last one referred to as height/length proportion in other studies using a classical morphometric approach (Ciocco, 1992; Gould, 1971; Orensanz, 1986). Both continuous characters have been related to the swimming capability of scallops (Gould, 1971; Stanley, 1972, 1988). While an elliptically elongated shell-disc (in the anteroposterior sense, with smaller height/length proportion; Fig. 2) is better to balance forces during the swimming making it easier, a more circular shell-disc shape (larger height/length proportion) has the opposite effect and can be associated with more sedentary habits (Gould, 1971). On the other hand, auricle symmetry is a feature associated with swimming activity, while an asymmetrically elongated anterior auricle may prevent the overturning of byssally attached individuals (Stanley, 1988, 1972). Under such theoretical frame, the symmetric auricles and the elliptically elongated shell-disc shape of scallops from LCO in San Matías Gulf suggest a more active swimming behavior than scallops from the San José Gulf and NEC location. This finding agrees with Orensanz (1986), who found a smaller height/length proportion in San Matías Gulf than in San José Gulf. Such shape was attributed to the high predation pressure in San Matías Gulf that may favor swimming escape responses. Instead, in San José Gulf, the wind-induced beach stranding exposure may be responsible of favoring a more sedentary shell shape (Orensanz, 1986). The fact that the morphologic differences between San Matías and San José gulfs in 1974-1976 (Orensanz, 1986) are similar to the finding in the present study suggest that the morphologic differentiation may have long-term stability. Further efforts should be directed to include more locations and distinct cohorts to test the temporal stability of the phenotypic trends (Cadrin, 2020).

Within San José Gulf, LES and PTE constituted a separated cluster (Figs 4, 6) with the most sedentary phenotypes (i.e., more circular shell-disc shape and asymmetrically small auricles). This morphologic differentiation is coincident with the presence of the San Jose front, as LES and PTE are located on each extreme of the front (Fig. 1). Larger morphologic differentiation at these places was previously noticed for the striped clam, *Ameghinomya antiqua* (Márquez et al., 2010b). The morphology of these locations might be reflecting how local strong tidal currents at these places drive the shell outline. The coastal geomorphology in conjunction with the tidal regimes (Tonini and Palma, 2017) produces strong tidal currents at LES and PTE (Picallo, 1980; Tonini and Palma, 2009), which are less stronger in other locations as Bengoa beach (eastward PSR) that respond mainly to the wind-stress (Moreira et al., 2009).

The morphometric differences founded along San Matías and San José gulfs may be the result of phenotypic plasticity driven by environmental conditions. In this geographical region, the lack of genetic differentiation indicates a plausible scenario of a common pool of larvae seeding the San Matías and San José gulfs (Getino Mamet et al., 2021). In this sense, given the shared larval pool genetically homogeneous, the morphological differentiation could be related to the local environmental conditions (e.g., currents) in which each cohort develops.

Finally, at the Buenos Aires shelf waters, although NEC presents a shape similar to the consensus, samples were correctly identified (Table 2) indicating that NEC can be differenced from the remaining locations inhabiting both gulfs. Such results are relevant considering the opportunistic nature of the fishery in this region. For example, during the fishing pulse of 2002, in the Buenos Aires shelf region, around 20,000 tonnes were fished in only three months before being wiped out by the bottom-trawl-based industrial Patagonian scallop fleet (Lasta and Campodónico, 2003; Soria et al., 2016). To give an idea of its importance, such captures represent more than the sum of fifty years of landings from San José Gulf, since the beginning of the activity. Currently, those beds lack fishing interest and it is uncertain if they have recovered after the 2002 fishing pulse. If such fishery reactivates, the morphometric differentiation will be useful information to properly distinguish captures from Buenos Aires.

The plasticity of the shell shape variation detected in this work has relevance in the context of fishery management. The scallop fishery from San José Gulf is managed as a single stock under a Total Allowable Catch system (Cinti et al., 2011; Orensanz and Seijo, 2013; Soria et al., 2016). However, the recruitment locations and depths in the gulf are rather heterogeneous, and therefore, fishers often propose a rotational/zoning scheme to focalize the fishing pressure in concordance with the stock assessment information (Fiorda and Parma, 2015; Soria et al., 2017). Additionally, phenotypic traits can be used as natural tags to the identification of the geographic origin of landings during the initial steps of the commercial process (Ibáñez, 2015, 2014; Shepard et al., 2010). The identification of the origin gives traceability to the capture, which is relevant to aid in law enforcement under a spatially structured management (Ortea and Gallardo, 2015). For example, the morphometric characterization of each bed could be relevant to control the origin of the scallops when some areas of the gulf are temporally closed due to a red tide event, which is a frequent phenomenon in the northern Patagonian gulfs (Sastre et al., 2018).

## 5. Conclusion

The findings of this study have relevance not only for the understanding of ecological adaptation to distinct environments of Tehuelche scallop but also for its fishery management. The shape differences between morphotypes complement the main distinction solely by the number of ribs and, together with the genetic differentiation, reinforce the idea of the presence of two ecotypes. Within the northern *tehuelchus* morphotype, the shell shape variation might be associated with environmental conditions indicating the plasticity of the shape outline. Further work is needed to understand the temporal stability of the phenotypic plasticity. Likewise, other biologic variables that may affect the swimming behavior (as predators, epibionts loads, bottom current, growth rate, dense-dependence) need to be studied. The findings of the present study have relevance for phenotypic stock identification that could be useful to know the origin of captures during the initial steps of the fishing process if zoning and rotational fishing schemes are implemented.

## CRediT authorship contribution statement

### Getino Mamet Leandro Nicolás

Conceptualization, Methodology, Formal analysis, Investigation, Visualization, Writing - Original Draft;

### Gaspar Soria

Conceptualization, Resources, Methodology, Formal analysis, Investigation, Funding acquisition, Project administration, Supervision, Writing - Review & Editing;

### Laura Schejter

Resources, Writing - Review & Editing;

### Federico Márquez

Resources, Methodology, Formal analysis, Writing - Review & Editing.

## Acknowledgments

We thank Dra. A. Parma for revising and providing helpful and insightful comments on this manuscript. We acknowledge Dr. E Morsán, P. Fiorda, J. Ascorti, R. Vera, F. Quiroga, T. Brochado, F. Irigoyen, N. Ortiz, J. Rúa, for assistance with data collection. Thanks to the Administración de Parques Nacionales (APN) and their staff at Reserva Natural de la Defensa Punta Buenos Aires for their assistance with the fieldwork. We collected Tehuelche scallop samples under permit DISPOSICION N 042 SsCyAP/14 to Ana Parma.

